# An accurate phasing tool via high-quality heterozygous variations

**DOI:** 10.64898/2025.12.21.695866

**Authors:** Jun Zhang, Fan Nie, Jianxin Wang

## Abstract

Long-read phasing has become a widely adopted approach for reconstructing paternal and maternal haplotypes in diploid genomes. However, most existing phasing methods treat all variants equally, even though low-quality variants frequently introduce switch errors. We present a new phasing method, snpphaser, which first phases high-quality variants and then leverages them to filter out low-quality variants that are inconsistent with the high-confidence haplotypes. We benchmarked snpphaser against state-of-the-art phasing methods across nine diverse datasets. The results demonstrate that snpphaser achieves superior contiguity and accuracy compared to existing approaches.

## 1 Introduction

The reconstruction of paternal and maternal haplotypes in diploid genomes is crucial for advancing our understanding of genetic variation, disease associations, and evolutionary processes (Tewhey *et al*., 2011). With short-read sequencing, haplotype phasing is inherently limited by read length and therefore often relies on supplementary information, such as linkage disequilibrium patterns in populations or Mendelian inheritance in pedigrees (Garg, 2021). In contrast, third-generation sequencing technologies (e.g., PacBio and Oxford Nanopore) generate substantially longer reads, which are increasingly used for human genome sequencing and assembly. Long-read-based phasing has thus emerged as the preferred strategy, as the extended read lengths enable the linkage of a greater number of alleles, thereby producing longer and more contiguous haplotype blocks.

Over the past decade, numerous read-based phasing methods have been developed. Among them, WhatsHap (Martin *et al*., 2016; Patterson *et al*., 2015) has become the most widely used tool across diverse applications. It operates on an allele matrix, where each column corresponds to a heterozygous variant and each row corresponds to a sequencing read. The algorithm partitions the reads into two conflict-free sets by solving the NP-hard weighted Minimum Error Correction (wMEC) problem. Although WhatsHap employs dynamic programming and down-samples to 15-fold coverage by default, phasing a human genome still requires several hours. HapCUT2 (Edge *et al*., 2017), another widely adopted method, assembles haplotypes from long reads using a likelihood-based max-cut procedure. However, it requires preprocessing of the input data and similarly takes several hours to phase a human genome. Margin (Ebler *et al*., 2019; Shafin *et al*., 2021) (formerly MarginPhase) uses a hidden Markov model to jointly perform genotype calling and haplotype phasing. It first selects a set of high-quality reads and variants for initial phasing and subsequently leverages these phased variants and reads to assign haplotypes to lower-quality variants and reads. Margin achieves human genome phasing within 1–2 hours while maintaining high accuracy and contiguity. LongPhase (Lin *et al*., 2022), in contrast, employs a greedy algorithm designed to maximize efficiency and contiguity. It is capable of phasing a human genome in only 10–20 minutes. However, we observed that LongPhase often produces switch errors, particularly in regions containing low-quality SNP calls or weak linkage between adjacent SNPs.

High-quality variants play a critical role in accurate haplotype phasing, whereas low-quality variants frequently introduce ambiguity and result in switch errors. To assess variant quality, we analyzed three sequencing datasets—Nanopore ultra-long, PacBio HiFi, and Nanopore R10.4 chemistry—from three samples (HG002, HG003, and HG004). SNPs were called using Clair3 (Zheng *et al*., 2022) and subsequently evaluated against the Genome in a Bottle (GIAB) (Zook *et al*., 2016) phased benchmark provided by NIST (v4.2.1). We observed that SNPs included in the benchmark were substantially enriched for high-quality variants, with proportions ranging from 83.55% to 85.88% for Nanopore ultra-long reads, 94.88% to 94.92% for PacBio HiFi reads, and 93.75% to 93.80% for Nanopore R10.4 reads (Supplementary Table S1). Moreover, in the more comprehensive phased benchmark of HG002, as many as 96.64% to 99.94% of SNPs are high-quality (Supplementary Table S2).

Here, we introduce a novel phasing method, snpphaser. The core strategy of snpphaser is to partition variants into high-quality and low-quality SNPs. In the first step, high-quality SNPs are phased using a greedy algorithm. Low-quality SNPs that are inconsistent with the high-confidence haplotypes are subsequently filtered out. The final phasing is then performed on the remaining set of both high-quality and retained low-quality SNPs. To further improve accuracy, the linkage between two adjacent SNPs is severed when the supporting evidence is weak—for example, when the number of reads connecting the two variants falls below a predefined threshold or when base quality scores are low at the corresponding positions. We benchmarked snpphaser against several state-of-the-art phasing tools, including WhatsHap, HapCUT2, Margin, and LongPhase. The results demonstrate that snpphaser achieves lower switch error rates while maintaining a comparable phased block N50.

## 2 Methods

Snpphaser partitions SNPs into two sets based on their quality annotations in the VCF file. Reads spanning at least two high-quality SNPs are first used to phase the high-quality variants. Leveraging these confidently phased SNPs, low-quality variants that are inconsistent with the high-quality haplotypes are filtered out. The final phasing is then performed on the combined set of high-quality SNPs and the remaining low-quality SNPs. To further improve accuracy, links between adjacent SNPs are removed when the supporting evidence is weak. Finally, snpphaser exploits the long-range information provided by long reads to connect adjacent phased blocks (Figure 1).

**Figure 1.**
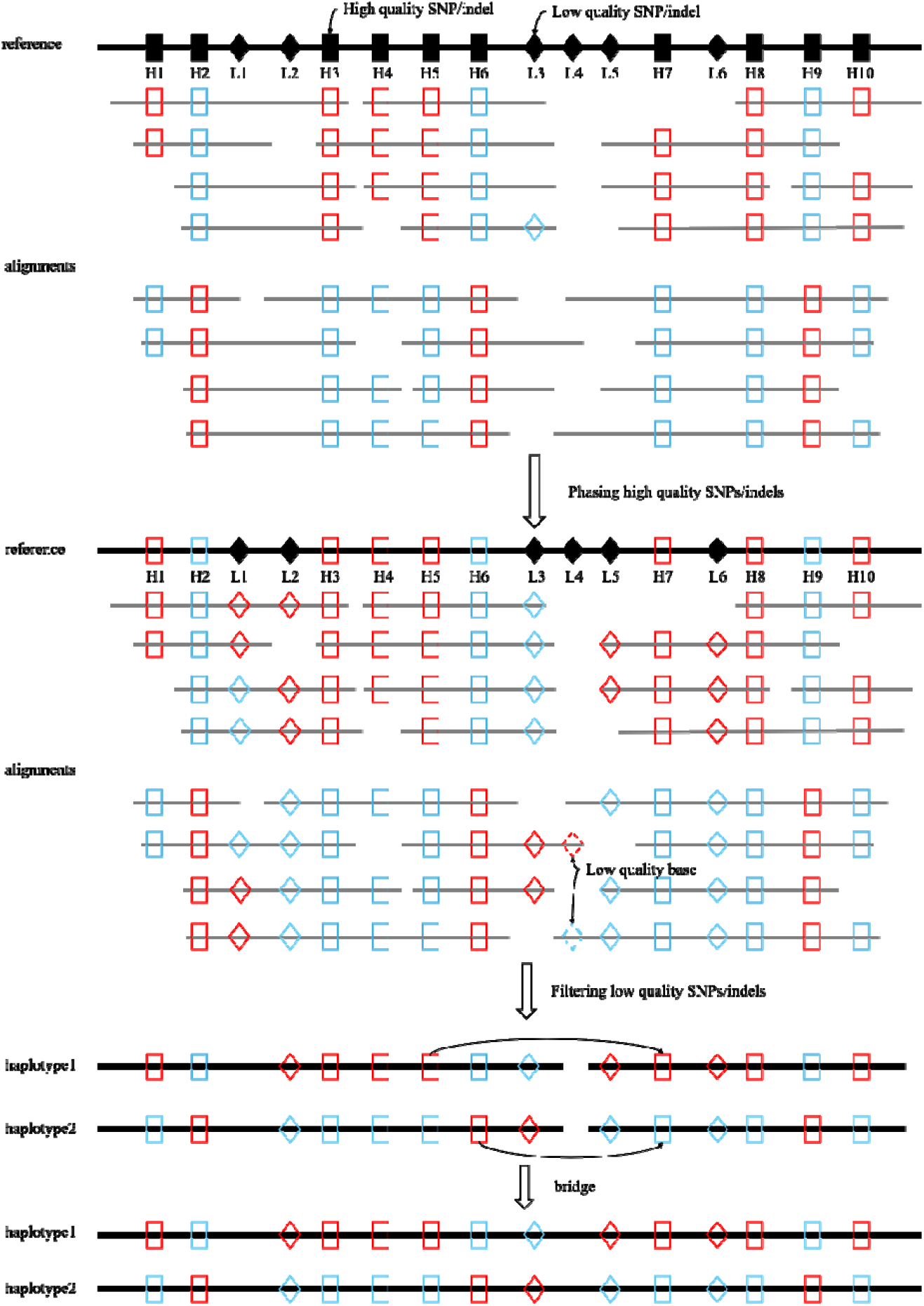
The workflow of snpphaser.

### 2.1 Dataset

We obtained Nanopore ultra-long and PacBio HiFi sequencing data for HG002, HG003, and HG004 from the Genome in a Bottle (GIAB) consortium, as well as Nanopore R10.4 chemistry data for the same samples from EPI2ME Labs. The nine datasets were aligned to the human reference genome (GRCh38) using minimap2 (Li, 2018, 2021), with parameters optimized for PacBio HiFi (-*x map* - *hifi*) and Nanopore (-*x map* - *ont*) reads. SNPs for each sample were subsequently called using Clair3. Across the datasets, more than three million heterozygous SNPs were identified, of which over 75% are high-quality (Supplementary Table S1).

### 2.2 Phasing of high-quality SNPs

Snpphaser incorporates the phasing algorithm of LongPhase to achieve efficient genome-wide haplotype assembly. The algorithm constructs a directed acyclic graph (DAG) in which vertices correspond to heterozygous variants ordered by genomic position. For each variant, two vertices (*v* and *v*′) are created to represent the reference (REF) and alternative (ALT) alleles. For every pair of adjacent variants, vi and vj, four directed edges are introduced: *e*(*v*_*i*_, *v*_*j*_), *e*(*v*_*i*_, *v*′_*j*_), *e*(*v*′_*i*_, *v*_*j*_), and *e*(*v*′_*i*_, *v*′_*j*_). Each edge is weighted by the number of sequencing reads supporting the linkage between the two alleles. The phasing problem is then formulated as identifying two disjoint paths through the graph that maximize the sum of edge weights, corresponding to the two haplotypes. Because of the bipartite structure and locality of the graph, this optimization can be solved efficiently using a greedy algorithm, enabling rapid genome-wide phasing.

For any two adjacent variants, only two mutually exclusive edge sets are possible: {*e*(*v*_*i*_,*v*_*j*_), *e*(*v*′_*i*_, *v*′_*j*_} or {*e*(*v*_*i*_, *v*′_*j*_), *e*(*v*′_*i*_, *v*_*j*_). The final haplotype paths must therefore be constructed from one of these two edge sets. Consequently, the optimal solution corresponds to selecting the set with the larger total edge weight. This property allows the problem to be solved efficiently using a greedy algorithm. Following the topological order of SNPs, the algorithm iteratively constructs disjoint paths by selecting, for each pair of adjacent variants, the edge set with the greater weight. The union of these locally optimal edge sets across all adjacent variant pairs yields the longest pair of disjoint haplotype paths. If no reads connect two adjacent variants, or if the weights of the two edge sets are equal, the paths are terminated at that position.

### 2.3 Filtering low-quality SNPs

Because low-quality SNPs frequently introduce switch errors, snpphaser selectively retains only high-confidence low-quality SNPs for phasing. Using the set of high-quality phased SNPs as a backbone, any low-quality SNP that is inconsistent with its neighboring high-quality variants is filtered out. For example, as illustrated in Figure 1, the low-quality SNP *L*1 is consistent with the high-quality SNP *H*1 but inconsistent with surrounding high-quality SNPs; therefore, it is removed. This procedure eliminates a large number of erroneous low-quality variants. After filtering, the remaining low-quality SNPs are combined with high-quality SNPs for the final phasing step, thereby improving overall performance. During this process, links between adjacent SNPs are discarded if their support is weak—specifically, when the number of reads spanning the two SNPs falls below a predefined threshold or when the base quality at the SNP positions is low. This constraint ensures higher phasing accuracy.

### 2.3 Recuse of entire long reads

When the linkage between two adjacent SNPs is weak, the corresponding edge is removed, which results in the fragmentation of a long haplotype block into two or more shorter blocks. To reconnect these fragmented blocks, snpphaser leverages the long-range linkage information inherent in long reads. Specifically, for each pair of adjacent phased blocks, snpphaser extracts all reads spanning both blocks. Each haplotype within the blocks is represented as a vertex, and edges correspond to reads spanning SNPs across the two haplotypes. Edge weights are defined as the number of SNPs within the reads that support a given haplotype connection. The bridging process is then resolved using the same greedy algorithm applied in the local phasing step. In the initial phasing stage, SNPs are phased solely based on short-range linkage between adjacent variants. This local strategy can be sensitive to sequencing errors in the absence of long-range linkage. To mitigate such errors, snpphaser further incorporates the full span of long reads to correct local phasing inconsistencies and improve overall haplotype accuracy.

### 2.4 Evaluation

To assess the phasing performance of each program, the phased results were evaluated against the GIAB phased benchmark provided by NIST (v4.2.1) using *WhatsHap campare*. The three benchmarks contained 454,285, 448,301, and 463,724 phased SNPs, respectively. Two standard accuracy metrics—switch error rate and block-wise Hamming distance—were computed with *WhatsHap campare*. In addition, an extended benchmark for HG002, generated using Strand-seq and trio-based phasing, was obtained from NIST (HG002_GRCh38_1_22_v4.2.1_benchmark_phased_MHCassembly_StrandSeqAND Trio.vcf.gz). This benchmark contains 1,755,615 phased SNPs.

## 3 Results

We compared snpphaser with four state-of-the-art phasing tools—WhatsHap, HapCUT2, Margin, and LongPhase—using three human genomes (HG002, HG003, and HG004) from the Genome in a Bottle (GIAB) consortium. The samples were sequenced with three technologies: Nanopore ultra-long PromethION reads (N50 = 43–48 kb), PacBio HiFi reads (CCS, 15 kb/20 kb chemistry2; N50 = 13–16 kb), and Nanopore R10.4 chemistry reads (N50 = 29–32 kb) (Supplementary Table S3). Raw reads were aligned to the human reference genome (GRCh38) using minimap2, with parameters optimized for PacBio HiFi (-*x map* - *hifi*) and Nanopore (-*x map* - *ont*) data. SNPs were called using Clair3, phased with the five programs, and evaluated against GIAB benchmarks using the *WhatsHap campare* command. In addition, we assessed performance on joint SNP/indel co-phasing.

### 3.1 Performance on Nanopore ultra-long datasets

On the Nanopore ultra-long datasets, the phased block N50s achieved by snpphaser (21 Mbp for HG002 and 17 Mbp for HG003) was comparable to those of LongPhase (21 Mbp for HG002 and 17 Mbp for HG003) and exceeded those produced by WhatsHap (18 Mbp for HG002 and 15 Mbp for HG003), Margin (13 Mbp for HG002 and 12 Mbp for HG003), and HapCUT2 (19 Mbp for HG002 and 15 Mbp for HG003) (Figure 2a; Supplementary Table S4). For HG004, LongPhase yielded the longest phased block N50 (24 Mbp), followed by snpphaser (20 Mbp), HapCUT2 (19 Mbp), WhatsHap (18 Mbp), and Margin (14 Mbp). With respect to accuracy, the switch error rates of snpphaser (0.22–0.26%) were slightly lower than those of WhatsHap (0.32–0.39%), Margin (0.39–0.43%), HapCUT2 (0.35–0.46%), and LongPhase (0.23–0.29%) across the three datasets (Figure 2b; Supplementary Table S4). Consistent results were observed when comparing block-wise Hamming distances. Margin phased the highest proportion of SNPs (>98%), followed by HapCUT2, WhatsHap, LongPhase, and snpphaser. In terms of runtime, snpphaser completed phasing in 14–34 minutes, whereas LongPhase required 8–23 minutes, Margin approximately 2 hours, and WhatsHap and HapCUT2 between 6 and 12 hours on the same datasets (Supplementary Table S4).

**Figure 2.**
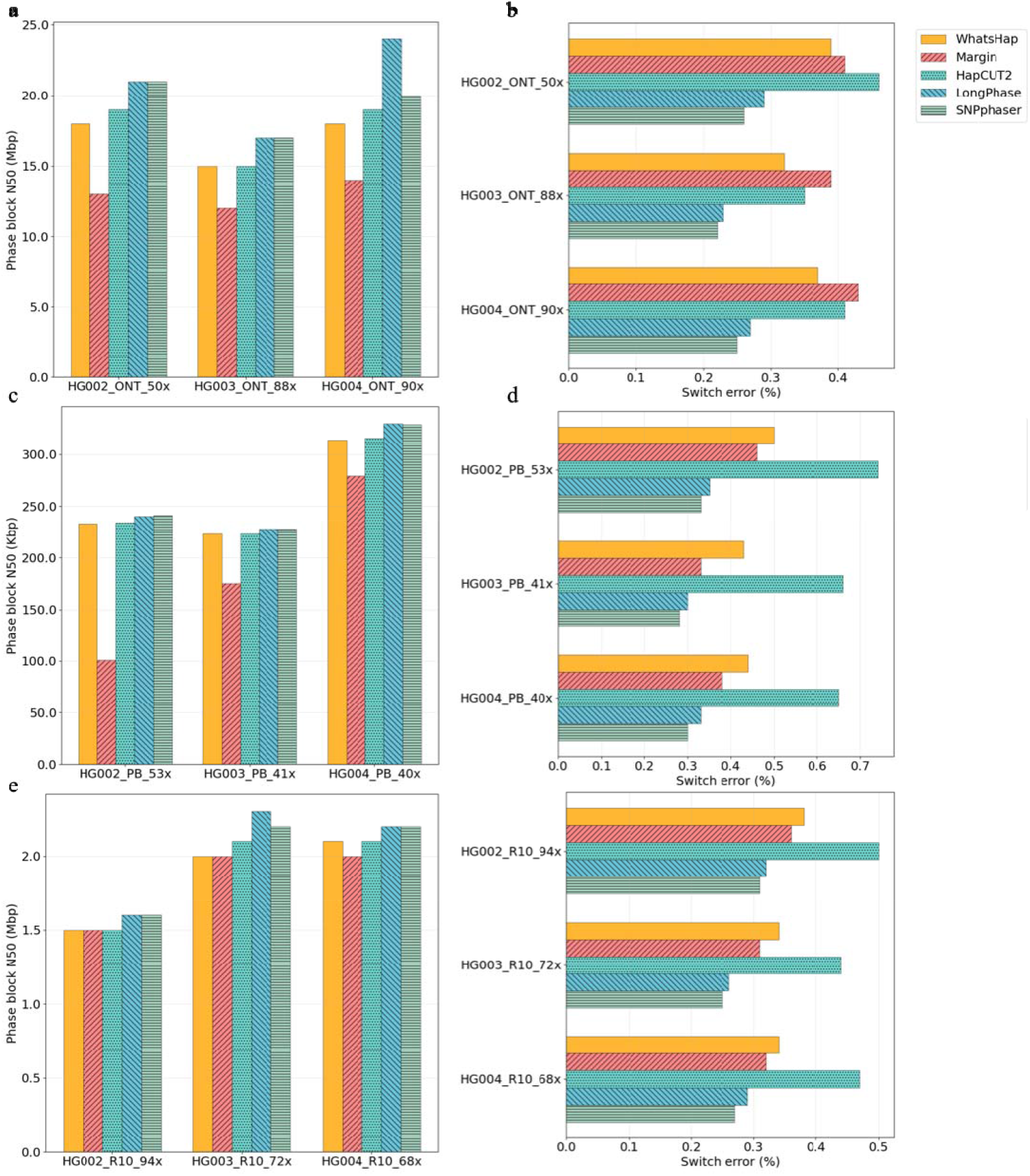
SNP-only phasing comparison of snpphaser, WhatsHap, Margin, HapCUT2 and LongPhase. (a) phasing contiguity on Nanopore ultra-long datasets, (b) switch error on Nanopore ultra-long datasets, (c) phasing contiguity on PacBio HiFi datasets, (d) switch error on PacBio HiFi datasets, (e) phasing contiguity on Nanopore R10 datasets, (f) switch error on Nanopore R10 datasets.

When co-phasing SNPs and indels, snpphaser and LongPhase both produced substantially longer phased blocks (84–87 Mbp) compared with WhatsHap (40–45 Mbp), HapCUT2 (40–46 Mbp), and Margin (16–17 Mbp) (Supplementary Figure S1a; Supplementary Table S5). The relative accuracy among the methods was consistent with that observed for SNP-only phasing (Supplementary Figure S1b; Supplementary Table S5). Specifically, snpphaser achieved the lowest switch error rate (0.21–0.26%) and block-wise Hamming distance, followed by LongPhase (0.23–0.30%), Margin (0.43–0.45%), HapCUT2 (0.48–0.59%), and WhatsHap (0.51–0.54%). In terms of runtime, snpphaser was slower than LongPhase but faster than Margin, HapCUT2, and WhatsHap (Supplementary Table S5).

### 3.2 Performance on PacBio HiFi datasets

On the PacBio HiFi datasets, the phased block N50s produced by snpphaser (228–329 kbp) were comparable to those of LongPhase (228–330 kbp), and both were slightly larger than those of WhatsHap (223–314 kbp), Margin (101–279 kbp), and HapCUT2 (223–316 kbp) (Figure 2c; Supplementary Table S4). Snpphaser achieved the lowest switch error rate and block-wise Hamming distance among the five programs, with values of 0.28–0.33%, compared with 0.43–0.50% for WhatsHap, 0.33–0.46% for Margin, 0.65–0.74% for HapCUT2, and 0.30–0.35% for LongPhase (Figure 2d; Supplementary Table S4). The percentage of phased SNPs was highest for LongPhase (>97%), followed by HapCUT2, WhatsHap, snpphaser, and Margin (Supplementary Table S4). In terms of computational efficiency, snpphaser and LongPhase completed phasing within 10–20 minutes, whereas WhatsHap and HapCUT2 required several hours (Supplementary Table S4).

For the co-phasing of SNPs and indels, snpphaser generated slightly shorter phased block N50s (330 kbp for HG002 and 314 kbp for HG003) than those produced by WhatsHap (333 kbp and 317 kbp, respectively) and HapCUT2 (332 kbp and 317 kbp, respectively), but longer than those of LongPhase (328 kbp and 313 kbp, respectively) and Margin (132 kbp and 262 kbp, respectively) (Supplementary Figure S1c; Supplementary Table S5). On the HG004 dataset, however, snpphaser achieved the longest phased block N50 (438 kbp), exceeding those of HapCUT2 (434 kbp), LongPhase (433 kbp), WhatsHap (432 kbp), and Margin (408 kbp). In terms of accuracy, snpphaser consistently yielded the lowest switch error rates (0.32–0.38%) as well as the smallest block-wise Hamming distances across all three datasets (Supplementary Figure S1d; Supplementary Table S5). Moreover, snpphaser required less runtime to complete phasing compared to WhatsHap, HapCUT2, and Margin (Supplementary Table S5).

### 3.3 Performance on Nanopore R10 datasets

We further evaluated the three most recent datasets sequenced with Nanopore R10.4 chemistry. On these datasets, the phased block N50s generated by LongPhase (1.6–2.3 Mbp) and snpphaser (1.6–2.2 Mbp) were comparable and slightly larger than those produced by WhatsHap (1.5–2.1 Mbp), Margin (1.5–2.0 Mbp), and HapCUT2 (1.5–2.1 Mbp) (Figure 2e; Supplementary Table S4). In terms of accuracy, snpphaser achieved the lowest switch error rates and block-wise Hamming distances, whereas HapCUT2 produced the highest (Supplementary Table S4). Regarding phasing completeness, Margin phased the largest proportion of SNPs, followed by LongPhase, WhatsHap, HapCUT2, and snpphaser (Supplementary Table S4). With respect to runtime, snpphaser required less computational time than Margin, WhatsHap, and HapCUT2, though it was slightly slower than LongPhase (Supplementary Table S4).

When performing co-phasing of SNPs and indels, WhatsHap produced the longest phased block N50s (7.5–10.3 Mbp), followed by HapCUT2 (7.6–10.3 Mbp), snpphaser (6.1–9.3 Mbp), LongPhase (5.1–7.8 Mbp), and Margin (4.0–4.7 Mbp) (Supplementary Figure S1e; Supplementary Table S5). In terms of accuracy, snpphaser achieved the lowest switch error rates across all three datasets (0.29–0.35%), consistent with the corresponding block-wise Hamming distance comparisons (Supplementary Figure S1f; Supplementary Table S5). The runtime comparison followed the same trend as observed in previous experiments (Supplementary Table S5).

We further down-sampled the Nanopore R10 datasets of HG002 to 60-fold and 30-fold coverage to assess the phasing performance of the five programs. The results are summarized in Table 1 and Supplementary Table S6. For SNP-only phasing, LongPhase produced the longest phased block N50s (1.2–1.5 Mbp), followed by snpphaser (1.1–1.4 Mbp), HapCUT2 (1.1–1.3 Mbp), WhatsHap (1.1–1.3 Mbp), and Margin (1.0–1.3 Mbp) (Table 1). In terms of accuracy, snpphaser achieved the lowest switch error rates and block-wise Hamming distances, whereas HapCUT2 produced the least accurate results. When evaluating SNP/indel co-phasing, HapCUT2 generated the longest phased block N50s (4.4–6.4 Mbp), followed by WhatsHap (4.3–6.3 Mbp), snpphaser (3.5–5.4 Mbp), LongPhase (3.3–4.7 Mbp), and Margin (2.8–3.2 Mbp). Consistent with the SNP-only results, snpphaser yielded the lowest switch error rates and block-wise Hamming distances.

**Table 1.** SNP-only phasing statistics on different coverage.

| Dataset | method | Phased pair | Switch error/% | Phased SNPs (%) | No. of block | Block N50 (bp) | Block-wise hamming distance/% | Time (min) |
| --- | --- | --- | --- | --- | --- | --- | --- | --- |
| 60X | WhatsHap | 116,094 | 460/0.40 | 96.10 | 4,527 | 1,318,790 | 408/0.35 | 244 |
|  | Margin | 114,805 | 416/0.36 | 98.24 | 4,269 | 1,317,820 | 371/0.32 | 49 |
|  | HapCUT2 | 116,015 | 586/0.51 | 96.61 | 4,461 | 1,334,067 | 527/0.45 | 337 |
|  | LongPhase | 112,526 | 369/0.33 | 95.35 | 3,873 | 1,463,774 | 327/0.29 | 14 |
|  | snpphaser | 114,808 | 353/0.31 | 93.61 | 3,606 | 1,424,180 | 313/0.27 | 22 |
| 30X | WhatsHap | 117,861 | 469/0.40 | 96.28 | 5,949 | 1,072,473 | 422/0.36 | 185 |
|  | Margin | 116,861 | 465/0.40 | 98.32 | 5,921 | 1,045,329 | 415/0.36 | 27 |
|  | HapCUT2 | 117,747 | 674/0.57 | 96.67 | 5,837 | 1,078,996 | 606/0.51 | 192 |
|  | LongPhase | 114,018 | 377/0.33 | 95.56 | 5,598 | 1,155,029 | 337/0.30 | 6 |
|  | snpphaser | 116,084 | 351/0.30 | 93.89 | 5,147 | 1,144,872 | 313/0.27 | 12 |

Since a more comprehensive benchmark is available for HG002, we downloaded it and used it as the ground truth to evaluate the phasing performance of the five programs. The results are presented in Supplementary Table S7. For SNP-only phasing, snpphaser achieved slightly lower switch error rates and block-wise Hamming distances compared with most tools, whereas Margin obtained the lowest values for both metrics.

## 4 Conclusions

Well-phased paternal and maternal haplotypes are fundamental for the analysis of genetic variation, disease associations, and evolutionary processes. Third-generation sequencing technologies, such as PacBio and Nanopore sequencing, enable haplotype phasing of SNPs owing to their long read lengths. Numerous tools have been developed for SNP phasing using long reads, including WhatsHap, HapCUT2, Margin, and LongPhase. Among these, our evaluations indicate that LongPhase generally produces longer phased block N50 values and higher accuracy. However, its performance is sensitive to low-quality SNPs and weak linkage between adjacent SNPs, which frequently lead to switch errors. To address this limitation, we developed a new phasing tool, snpphaser, which first phases high-quality SNPs and then uses the resulting haplotypes to filter out low-quality SNPs that are inconsistent with the high-quality set. Our evaluation demonstrates that snpphaser achieves more accurate phasing results while maintaining contiguity comparable to that of LongPhase.

## Acknowledgements

This work was supported in part by the National Natural Science Foundation of China Grant (Nos. 62350004, 62332020), and the Project of Xiangjiang Laboratory (No. 23XJ01011).

## Supplementary material for

**Supplementary Figure S1.**
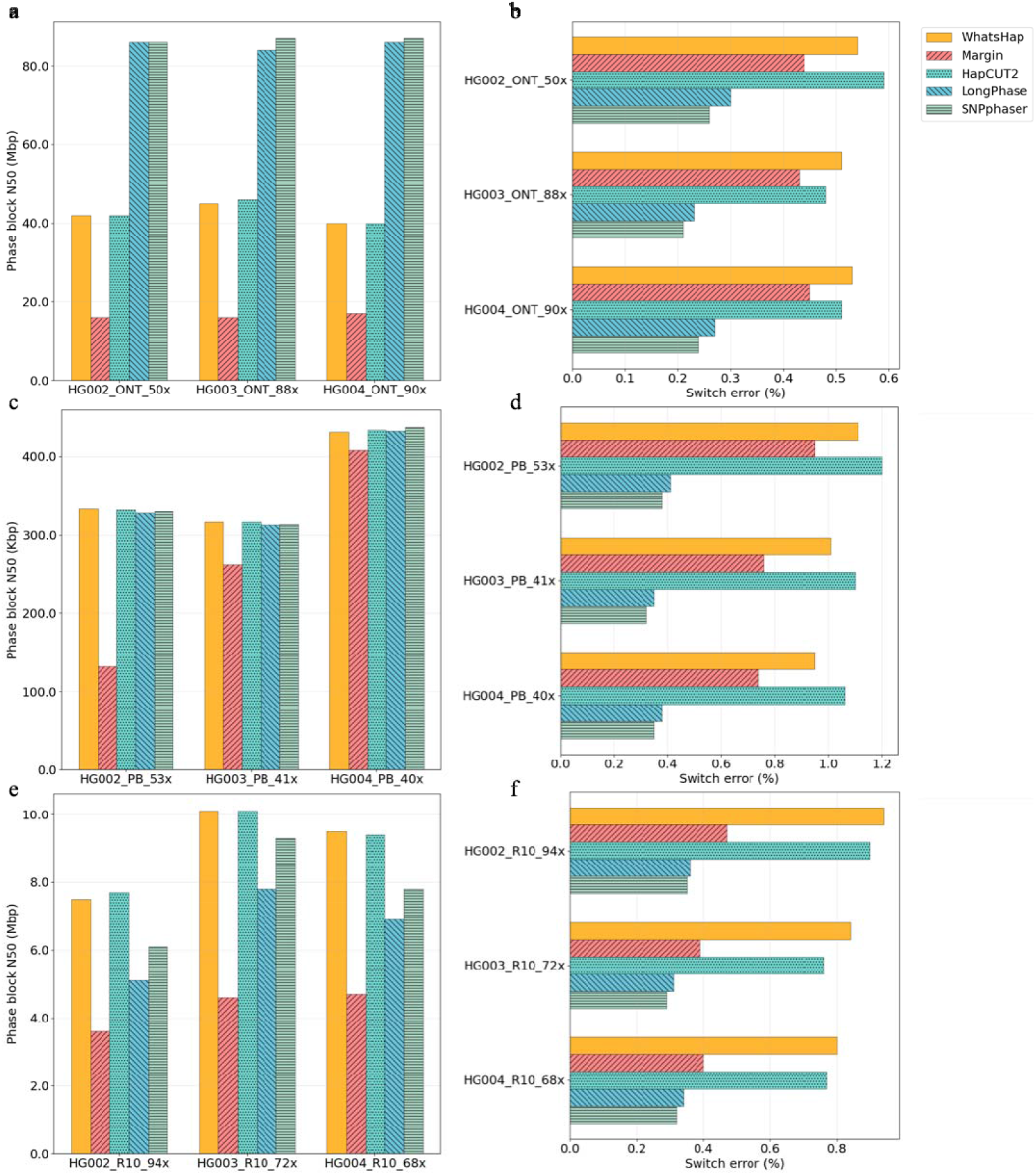
SNP/indel co-phasing comparison of snpphaser, WhatsHap, Margin, HapCUT2 and LongPhase. (a) phasing contiguity on Nanopore ultra-long datasets, (b) switch error on Nanopore ultra-long datasets, (c) phasing contiguity on PacBio HiFi datasets, (d) switch error on PacBio HiFi datasets, (e) phasing contiguity on Nanopore R10 datasets, (f) switch error on Nanopore R10 datasets.

**Supplementary Table S1.** SNP called statistics of HG002, HG003, and HG004 using GIAB phased benchmark in NIST (v4.2.1)

| sequence | sample | Homozygous | Heterozygous | High quality | Heterozygous | High quality |
| --- | --- | --- | --- | --- | --- | --- |
|  |  | SNP | SNP | Heterozygous | SNP phased in | heterozygous SNP |
|  |  |  |  | SNP | GIAB | phased in GIAB/% |
| Nanopore<br>ultra-long | HG002 | 3,618,619 | 3,136,581 | 2,370,028 | 281,539 | 235,223/83.55 |
|  | HG003 | 3,701,902 | 3,095,998 | 2,388,457 | 280,892 | 241,234/85.88 |
|  | HG004 | 3,725,166 | 3,142,736 | 2,497,688 | 289,116 | 248,168/85.84 |
| PacBio<br>HiFi | HG002 | 2,031,716 | 3,222,020 | 2,932,863 | 313,787 | 297,831/94.92 |
|  | HG003 | 2,046,648 | 3,184,628 | 2,887,488 | 309,603 | 293,866/94.92 |
|  | HG004 | 1,973,435 | 3,328,336 | 3,029,087 | 319,608 | 303,236/94.88 |
| Nanopore<br>R10.4 | HG002 | 2,228,747 | 3,274,185 | 2,821,985 | 309,343 | 290,009/93.75 |
|  | HG003 | 2,201,287 | 3,230,855 | 2,782,164 | 304,886 | 285,994/93.80 |
|  | HG004 | 2,136,368 | 3,342,654 | 2,906,730 | 314,132 | 294,653/93.80 |

**Supplementary Table S2.** SNP called statistics of HG002 using GIAB phased benchmark in NIST (v4.2.1 MHCassembly_StrandSeqANDTrio)

| sequence | Homozygous | Heterozygous | High quality | Heterozygous | High quality |
| --- | --- | --- | --- | --- | --- |
|  | s SNP | us SNP | Heterozygous | SNP phased in | heterozygous SNP |
|  |  |  | SNP | GIAB | phased in GIAB/% |
| Nanopore ultra-long | 3,618,619 | 3,136,581 | 2,370,028 | 1,617,255 | 1,562,975/96.64 |
| PacBio HiFi | 2,031,716 | 3,222,020 | 2,932,863 | 1,754,489 | 1,753,412/99.94 |
| Nanopore R10.4 | 2,228,747 | 3,274,185 | 2,821,985 | 1,736,345 | 1,723,274/99.25 |

**Supplementary Table S3.** statistics of raw sequences.

| sequence | sample | Error rate (%) | No. of read | N50 | Total base | coverage |
| --- | --- | --- | --- | --- | --- | --- |
| Nanopore<br>ultra-long | HG002 | 8.58 | 19,328,993 | 48,393 | 156,533,436,510 | 50 |
|  | HG003 | 7.65 | 23,954,632 | 43,303 | 272,567,173,779 | 88 |
|  | HG004 | 8.19 | 29,319,334 | 46,689 | 279,586,438,438 | 90 |
| PacBio HiFi | HG002 | 0.69 | 11,714,594 | 13,593 | 166,222,697,393 | 53 |
|  | HG003 | 0.70 | 8,425,983 | 14,977 | 129,517,546,663 | 41 |
|  | HG004 | 0.66 | 7,180,342 | 16,796 | 123,812,043,382 | 40 |
| Nanopore<br>R10.4 | HG002 | 2.47 | 16,252,223 | 29,433 | 291,940,137,586 | 94 |
|  | HG003 | 1.95 | 16,507,232 | 32,907 | 224,226,429,038 | 72 |
|  | HG004 | 1.94 | 16,006,628 | 32,203 | 211,147,159,417 | 68 |

**Supplementary Table S4.**
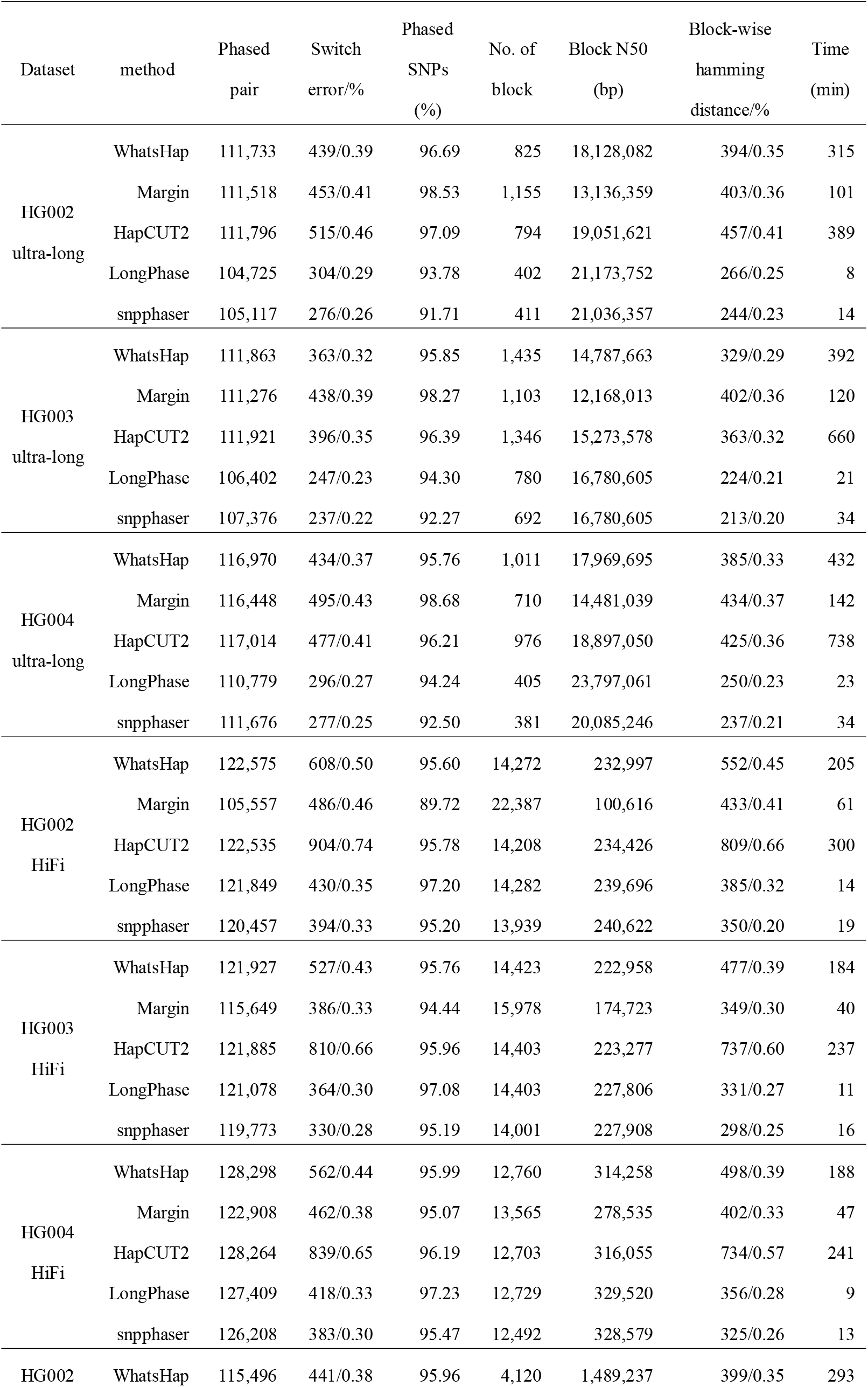
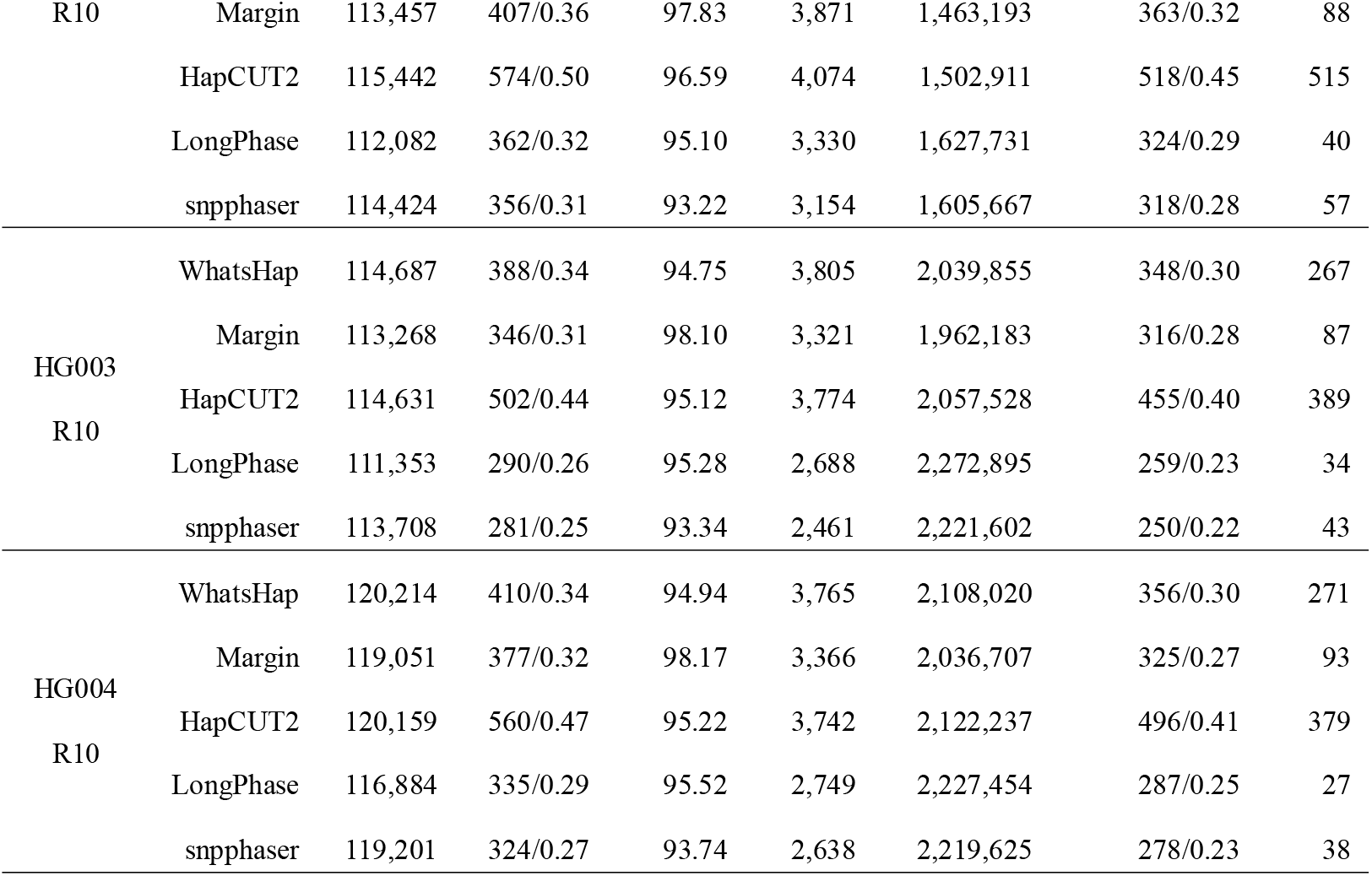
SNP-only phasing statistics.

**Supplementary Table S5.**
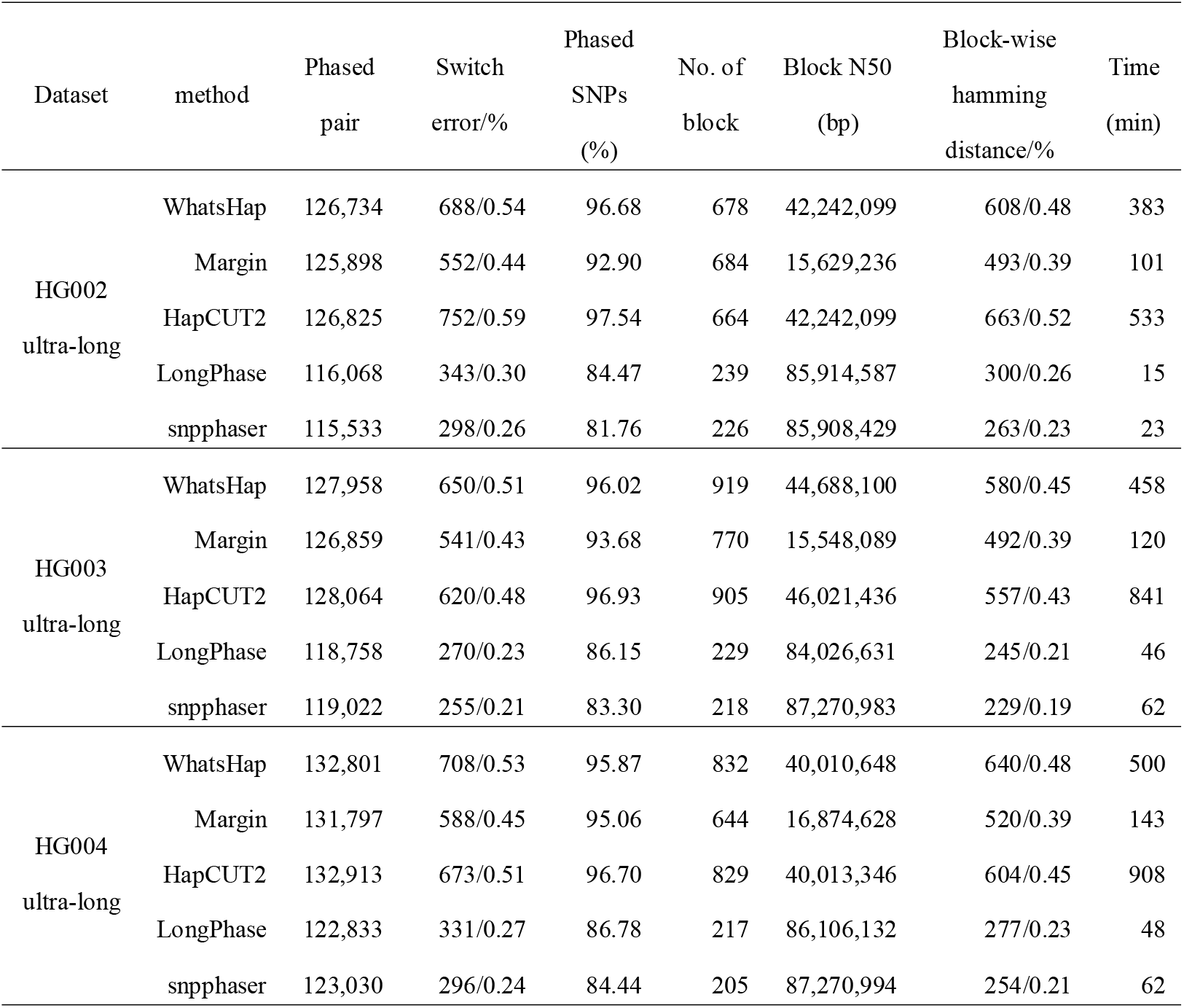
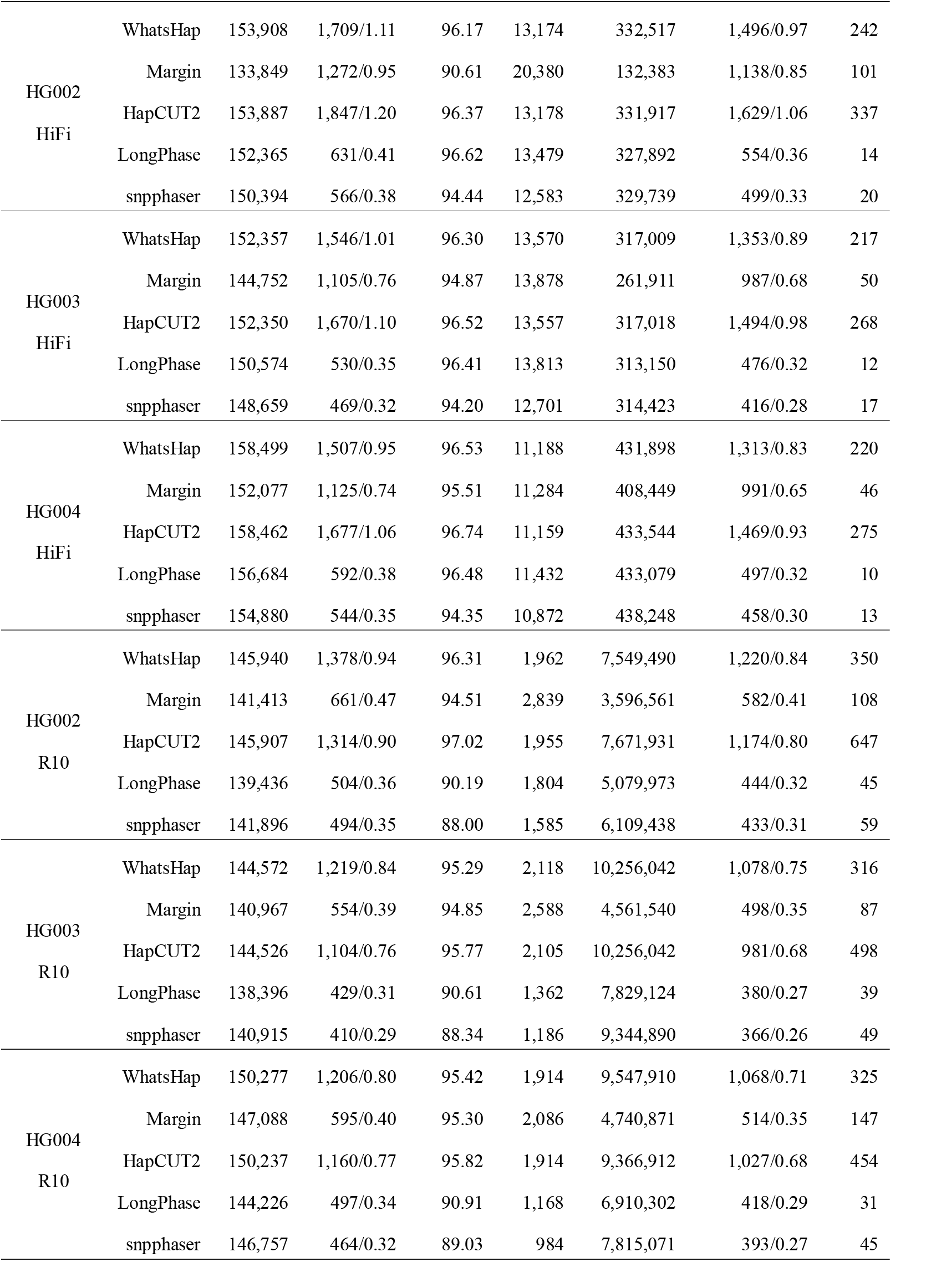
SNP/indel co-phasing statistics.

**Supplementary Table S6.** SNP/indel co-phasing statistics on different coverage.

| Dataset | method | Phased<br>pair | Switch<br>error/% | Phased<br>SNPs<br>(%) | No. of<br>block | Block N50<br>(bp) | Block-wise<br>hamming<br>distance/% | Time<br>(min) |
| --- | --- | --- | --- | --- | --- | --- | --- | --- |
| 60X | WhatsHap | 146,594 | 1,420/0.97 | 96.44 | 1,922 | 6,296,129 | 408/0.35 | 286 |
|  | Margin | 143,011 | 676/0.47 | 94.88 | 2,830 | 3,231,369 | 371/0.32 | 49 |
|  | HapCUT2 | 146,547 | 1,327/0.91 | 97.05 | 1,913 | 6,437,619 | 527/0.45 | 454 |
|  | LongPhase | 139,845 | 508/0.36 | 90.20 | 1,818 | 4,654,353 | 327/0.29 | 15 |
|  | snpphaser | 142,346 | 480/0.34 | 88.06 | 1,624 | 5,363,398 | 313/0.27 | 27 |
| 30X | WhatsHap | 148,151 | 1,355/0.91 | 96.55 | 1,883 | 4,292,510 | 422/0.36 | 227 |
|  | Margin | 145,177 | 785/0.54 | 94.21 | 2,618 | 2,835,912 | 415/0.36 | 27 |
|  | HapCUT2 | 148,020 | 1,463/0.99 | 97.16 | 1,863 | 4,387,480 | 606/0.51 | 237 |
|  | LongPhase | 140,984 | 522/0.37 | 89.00 | 1,937 | 3,334,084 | 337/0.30 | 8 |
|  | snpphaser | 143,406 | 485/0.34 | 86.57 | 1,798 | 3,528,874 | 313/0.27 | 15 |

**Supplementary Table S7.** Phasing statistics comparison on MHCassembly_StrandSeqANDTrio.

MHCassembly\_StrandSeqANDTrio
|  |  | SNP-only phasing |  |  | SNP/indel co-phasing |  |  |
| --- | --- | --- | --- | --- | --- | --- | --- |
|  | method | Phased<br>pairs | Switch<br>error/% | Block-wise<br>hamming<br>distance/% | Phased<br>pairs | Switch<br>error/% | Block-wise<br>hamming<br>distance/% |
| Ultra-long | Whatshap | 1,512,884 | 716/0.05 | 164,702/10.89 | 1,602,680 | 2,590/0.16 | 525,622/32.80 |
|  | Margin | 1,512,471 | 520/0.03 | 121,609/8.04 | 1,600,168 | 1,025/0.06 | 357,873/22.36 |
|  | Hapcut2 | 1,512,962 | 923/0.06 | 153,821/10.17 | 1,602,910 | 1,805/0.11 | 475,740/29.68 |
|  | Longphase | 1,492,181 | 575/0.04 | 212,678/14.25 | 1,562,059 | 968/0.06 | 556,090/35.60 |
|  | snpphaser | 1,488,895 | 568/0.04 | 204,946/13.76 | 1,555,790 | 766/0.05 | 476,046/30.60 |
| HiFi | Whatshap | 1,523,029 | 724/0.05 | 8,369/0.55 | 1,742,169 | 2,363/0.14 | 31,368/1.80 |
|  | Margin | 1,443,582 | 654/0.05 | 4,911/0.34 | 1,659,350 | 4,437/0.27 | 49,231/2.97 |
|  | Hapcut2 | 1,523,082 | 836/0.05 | 10,971/0.72 | 1,742,182 | 2,407/0.14 | 27,771/1.59 |
|  | Longphase | 1,522,185 | 713/0.05 | 12,741/0.84 | 1,738,586 | 1,992/0.11 | 29,991/1.73 |
|  | snpphaser | 1,522,723 | 666/0.04 | 7,997/0.53 | 1,740,062 | 1,908/0.11 | 25,007/1.44 |
| R10 | Whatshap | 1,533,435 | 665/0.04 | 28,775/1.88 | 1,726,815 | 7,293/0.42 | 430,838/24.95 |
|  | Margin | 1,533,038 | 669/0.04 | 35,643/2.32 | 1,707,972 | 2,494/0.15 | 243,465/14.25 |
|  | Hapcut2 | 1,533,457 | 783/0.05 | 31,158/2.03 | 1,727,345 | 3,516/0.20 | 433,054/25.07 |
|  | Longphase | 1,531,872 | 677/0.04 | 34,705/2.27 | 1,692,792 | 2,169/0.13 | 318,041/18.79 |
|  | snpphaser | 1,532,436 | 638/0.04 | 29,036/1.89 | 1,695,111 | 2,222/0.13 | 335,169/19.77 |
| R10 60X | Whatshap | 1,533,123 | 691/0.05 | 31,014/2.02 | 1,724,957 | 6,620/0.38 | 420,394/24.37 |
|  | Margin | 1,532,862 | 656/0.04 | 29,265/1.91 | 1,708,504 | 2,466/0.14 | 226,824/13.28 |
|  | Hapcut2 | 1,533,155 | 815/0.05 | 30,837/2.01 | 1,725,365 | 3,932/0.23 | 416,238/24.12 |
|  | Longphase | 1,531,424 | 699/0.05 | 38,591/2.52 | 1,691,734 | 2,408/0.14 | 338,635/20.02 |
|  | snpphaser | 1,531,892 | 644/0.04 | 26,495/1.73 | 1,694,498 | 2,414/0.14 | 361,634/21.34 |
| R10 30X | Whatshap | 1,530,366 | 821/0.05 | 40,874/2.67 | 1,716,777 | 5,926/0.35 | 380,437/22.16 |
|  | Margin | 1,530,058 | 779/0.05 | 39,167/2.56 | 1,704,775 | 3,232/0.19 | 246,765/14.47 |
|  | Hapcut2 | 1,530,382 | 994/0.06 | 41,464/2.71 | 1,716,831 | 5,244/0.31 | 355,610/20.71 |
|  | Longphase | 1,527,906 | 761/0.05 | 38,158/2.50 | 1,686,018 | 2,752/0.16 | 312,535/18.54 |
|  | snpphaser | 1,528,136 | 719/0.05 | 30,737/2.01 | 1,688,518 | 2,692/0.16 | 313,062/18.54 |

